# Effects of aluminum-salt, CpG and emulsion adjuvants on the stability and immunogenicity of a virus-like particle displaying the SARS-CoV-2 receptor-binding domain (RBD)

**DOI:** 10.1101/2023.07.10.548406

**Authors:** Ozan S. Kumru, Sakshi Bajoria, Kawaljit Kaur, John M. Hickey, Greta Van Slyke, Jennifer Doering, Katherine Berman, Charles Richardson, Hans Lien, Harry Kleanthous, Nicholas J. Mantis, Sangeeta B. Joshi, David B. Volkin

**Affiliations:** Department of Pharmaceutical Chemistry, Vaccine Analytics and Formulation Center, University of Kansas, Lawrence, KS 66047, USA; Division of Infectious Disease, Wadsworth Center, New York State Department of Health, Albany, NY 12208, USA; Icosavax, 1930 Boren Ave, Suite 1000. Seattle, WA 98101, USA; Bill and Melinda Gates Foundation, Seattle, WA 98109, USA

**Keywords:** Vaccine, Formulation, Stability, Aluminum-salt, CpG, Emulsion, Adjuvant, SARS-CoV-2, Antigen, VLP (Virus-Like Particle)

## Abstract

Second-generation COVID-19 vaccines with improved immunogenicity (e.g., breadth, duration) and availability (e.g., lower costs, refrigerator stable) are needed to enhance global coverage. In this work, we formulated a clinical-stage SARS-CoV-2 receptor binding domain (RBD) virus-like particle (VLP) vaccine candidate (IVX-411) with widely available adjuvants. Specifically, we assessed the *in vitro* storage stability and *in vivo* mouse immunogenicity of IVX-411 formulated with aluminum-salt adjuvants (Alhydrogel™, AH and Adjuphos™, AP), without or with the TLR-9 agonist CpG-1018™ (CpG), and compared these profiles to IVX-411 adjuvanted with an oil-in-water nano-emulsion (AddaVax™, AV). Although IVX-411 bound both AH and AP, lower binding strength of antigen to AP was observed by Langmuir binding isotherms. Interestingly, AH- and AP-adsorbed IVX-411 had similar storage stability profiles as measured by antigen binding assays (competitive ELISAs), but the latter displayed higher pseudovirus neutralizing titers (pNT) in mice, at levels comparable to titers elicited by AV-adjuvanted IVX-411. CpG addition to alum (AP or AH) resulted in a marginal trend of improved pNTs in stressed samples only, yet did not impact the storage stability profiles of IVX-411. In contrast, previous work with AH-formulations of a monomeric RBD antigen showed greatly improved immunogenicity and decreased stability upon CpG addition to alum. At elevated temperatures (25, 37°C), IVX-411 formulated with AH or AP displayed decreased *in vitro* stability compared to AV-formulated IVX-411and this rank-ordering correlated with *in vivo* performance (mouse pNT values). This case study highlights the importance of optimizing antigen-adjuvant interactions to develop low cost, aluminum-salt adjuvanted recombinant subunit vaccine candidates.

## Introduction

To date, the COVID-19 pandemic has resulted in >760 million confirmed cases and >6.9 million deaths worldwide ^1^. Although first-generation COVID-19 vaccines are currently available based on mRNA-LNP, adenoviral-vector, and subunit protein vaccine platforms ^2, 3^, there remains an urgent need for second-generation vaccines with improved immunogenicity to ensure efficacy against circulating variants of concern ^4, 5^ as well as greater global access. For example, mosaic nanoparticles with multimeric display of SARS-CoV-2 RBDs have been shown to induce a wide breadth of protection against many different SARS-like betacoronaviruses ^6, 7^. Such subunit vaccine candidates are well-suited for improving global access due to low cost-of-goods, high production capacity, and storage stability profiles compatible with refrigerated cold-chain infrastructure ^8–10^.

Subunit vaccines, however, generally require one or more immunostimulatory adjuvants to be effective. Several diverse adjuvant systems are available, which range from colloidal suspensions of aluminum-salts of varying chemical composition and morphology (generally referred to as alum), to oil-in-water emulsions, to specific immunostimulatory molecules (i.e., TLR agonists including oligonucleotides, phosphoryl lipid A, saponins) as well as combinations thereof ^11^. Although the choice of adjuvant depends foremost on clinical performance, additional practical considerations including cost, availability of GMP materials, and effects on antigen compatibility must be considered ^11^. This is especially true if the ultimate goal is to produce effective, lower-cost vaccines targeted for global use in low-and middle-income countries.

In this work, we focused on a subset of widely available adjuvants that are currently found in licensed vaccine products including aluminum-salts (alum), cytosine phosphoguanine (CpG) oligodeoxynucleotides and oil-in-water emulsions. These adjuvants differ in their composition, immunological effects, commercial availability, and cost. Alum adjuvants have been commonly used in vaccines for nearly a century, are inexpensive, readily available at large-scale, and have an excellent safety profile in a variety of infant, pediatric and adult vaccines ^12, 13^. The CpG 1018™ oligonucleotide adjuvant (referred to herein as CpG) is a TLR9 agonist with demonstrated safety and efficacy in subunit vaccines including a licensed hepatitis B vaccine (HEPLISAV-B®) as well as when co-formulated with alum in a SARS-CoV-2 RBD-based vaccine ^14–18^. The oil-in-water emulsion adjuvants MF59 and AS03 are currently licensed for use in seasonal and pandemic influenza vaccines (e.g., Fluad®, Focetria®, Pandemrix®, and Celura®). The major components of MF59 are squalene oil and the non-ionic surfactants Polysorbate 80 and Span 85 ^19, 20^. There are considerable costs associated with access to and manufacturing of oil-in-water emulsion adjuvants to GMP standards at large scale ^21, 22^. In addition, squalene is harvested from the livers of deep-sea sharks, which raises environmental, sustainability, and ecological concerns ^23^. Since alum and CpG adjuvants do not have these same concerns, this work with a second-generation COVID-vaccine antigen focused on comparing optimized formulations of alum (AH, AP), with and without CpG, as bench-marked to an oil-in-water emulsion adjuvant, AddaVax^TM^ (AV), which is equivalent to MF59, but unavailable for human use.

There are several different protein-based COVID-19 subunit vaccine candidates approved for use or under clinical development based on multimeric display of antigens on nanoparticles ^18, 24^. Multivalent presentation of antigens has been successfully implemented previously in commercial vaccines using virus-like particles (VLP), including licensed vaccines against Hepatitis B Virus (HBV) and Human Papilloma Virus (HPV). These VLP based vaccines consist of a viral surface protein (i.e., HBsAg and L1, for HBV and HPV, respectively), which self-assemble into VLPs that enable multimeric antigen display ^25, 26^. In general, multivalent presentation of an antigen on the surface of a VLP generates higher cellular and humoral immune responses when compared to monomeric antigens ^27–34^. Since VLPs are typically >10 fold larger by molecular weight when compared monomeric proteins angiens, VLPs are more efficiently taken up by antigen presenting cells (APC), proteotically processed, and presented by MHC class II. Furthermore, the repetitive conformational eptiopes on the surface of VLPs promotes crosslinking of surface immunoglobulin B-cell receptors, which leads a robust B-cell response ^35^.

One promising example of a COVID-19 vaccine antigen designed for multimeric antigen display on a nanoparticle is RBD-NPs, which are composed of a computationally designed nanoparticle (I53-50) engineered to display 60 copies of the Wuhan-1 SARS-CoV-2 receptor binding domain (RBD) ^31, 36^. RBD-NPs elicit robust immune responses in mice and non-human primates (NHPs) when formulated with adjuvants such as AH, AH+CpG, AS37, a TLR7 agonist adsorbed to alum, and various oil-in-water emulsions. RBD-NPs formulated with the AS03 and AH+CpG adjuvant systems provided protective immunity based on induction of virus neutralizing antibody titers ^31, 37, 38^. Furthermore, results from a Phase 1/2 and Phase 3 clinical trials ^39, 40^ demonstrated an RBD-nanoparticle-based vaccine adjuvanted with AS03 (GP150) was highly immunogenic and well-tolerated in humans and has now been approved for use in South Korea (SKYCovine® from SK Bioscience).

The major objective of this work was to assess the feasibility of developing a lower-cost, aluminum-salt adjuvanted formulation of a SARS-CoV-2 vaccine candidate composed of a two-component I53-50 nanoparticle displaying 60 copies of the SARS-CoV-2 Wuhan-1 RBD (termed IVX-411). Previous work has demonstrated that AV-adjuvanted IVX-411 induces robust neutralizing antibody responses in mice against both Wuhan-1 and variants of concern and protected Syrian hamsters in a SARS-CoV-2 challenge study ^29^. In this work, we first performed physicochemical characterization and antigen-adjuvant interaction studies of IVX-411 formulated with aluminum-salt, CpG and oil-in-water emulsion adjuvants. We then developed stability-indicating antigen binding assays to monitor the structural integrity of key epitopes of formulated IVX-411 in the presence of different adjuvants. Finally, we evaluated the storage stability and *in vivo* mouse immunogenicity profiles of the various IVX-411-adjuvant formulations. The results of this work are compared to previous work with similar formulations of monomeric RBD, and are discussed more generally in the context of formulation development of low-cost vaccine candidates targeted for use in low and middle income countries (LMICs).

## Materials and Methods

### Materials

IVX-411 antigen was provided frozen by Icosavax on dry ice at a protein concentration of 0.68 mg/mL in a formulation buffer (50 mM Tris, 150 mM NaCl, 0.1 M L-Arginine, 5% Sucrose, pH 8.0), thawed, aliquoted and stored at -80°C. CpG adjuvant was provided by Dynavax in a lyophilized powder form and was reconstituted in 20 mM Tris, 100 mM NaCl, pH 7.5 and stored at -20°C at a final concentration of 12-15 mg/mL. AH, AP, and AV adjuvants were purchased from InvivoGen (San Diego, CA). All other chemicals were of high purity and purchased from Sigma-EMD Millipore (Burlington, MA) except for sucrose, which was purchased from Pfanstiehl (Waukegan, IL).

### Methods

The analytical methods employed in this study, including physicochemical characterization techniques (i.e., SDS-PAGE, fourier transform infrared (FTIR) and intrinsic fluorescence (IF) spectroscopy, differential scanning calorimetry (DSC), sedimentation velocity analytical ultracentrifugation (SV-AUC), and dynamic light scattering (DLS), as well as antigen binding *in vitro* potency assays (competitive ELISAs) and mouse immunogenicity studies including pseudovirus neutralization titers are described in detail in the Supplemental Methods. Experimental procedures for antigen-adjuvant interaction (i.e., Langmuir adsorption isotherms) and storage stability studies are also provided in the Supplemental Methods. Briefly, for biophysical measurements in solution, IVX-411 was diluted to 0.1-0.3 mg/mL in the formulation buffer. For antigen-adjuvant binding studies, IVX-411 was diluted to 0.05-0.2 mg/mL in formulation buffer. For storage stability and mouse immunogenicity studies, IVX-411 was diluted to 2 mcg/mL in the formulation buffer with adjuvant concentrations of (i) 1.5 mg/mL AH, (ii) 1.5 mg/mL AP, (iii) 1.5 mg/mL AH and 0.3 mg/mL CpG, (iv) 1.5 mg/mL AP and 0.3 mg/mL CpG, or (v) 1X final concentration of AV.

## Results

### Analytical characterization of IVX-411 antigen

IVX-411 is a computationally designed, two-component VLP-based antigen consisting of the receptor binding domain (RBD) of the spike protein from SARS-CoV-2 genetically fused to a homotrimeric complex (Component A) and a homopentameric complex (Component B). When Components A and B are mixed in solution, a VLP (IVX-411) spontaneously self assembles into a nanoparticle with multimeric display of RBD, as represented schematically in **Figure 1A** ^31^. SDS-PAGE analysis (**Figure 1B**) shows Components A and B migrate at the expected ∼50 and ∼15 kDa molecular weight (MW) values, respectively. Higher MW species (∼90 kDa) were also observed in the non-reduced sample that were absent under reducing conditions, indicating the presence of disulfide-linked oligomers. After PNGase-F treatment, Component A shifted to a lower MW indicating the presence of N-linked glycans. An additional minor band at ∼20-25 kDa could represent the core trimeric domain of Component A without RBD. The diameter of IVX-411 was 28 ± 3 nm as determined by negative staining Transmission Electron Microscopy (TEM) (**Figure 1C**). TEM analysis of IVX-411 formulated with AH showed the antigen adsorbed to the adjuvant surface (**Figure 1D**). Similar results were observed when IVX-411 was adsorbed to AP adjuvant (data not shown).

**Figure 1.**
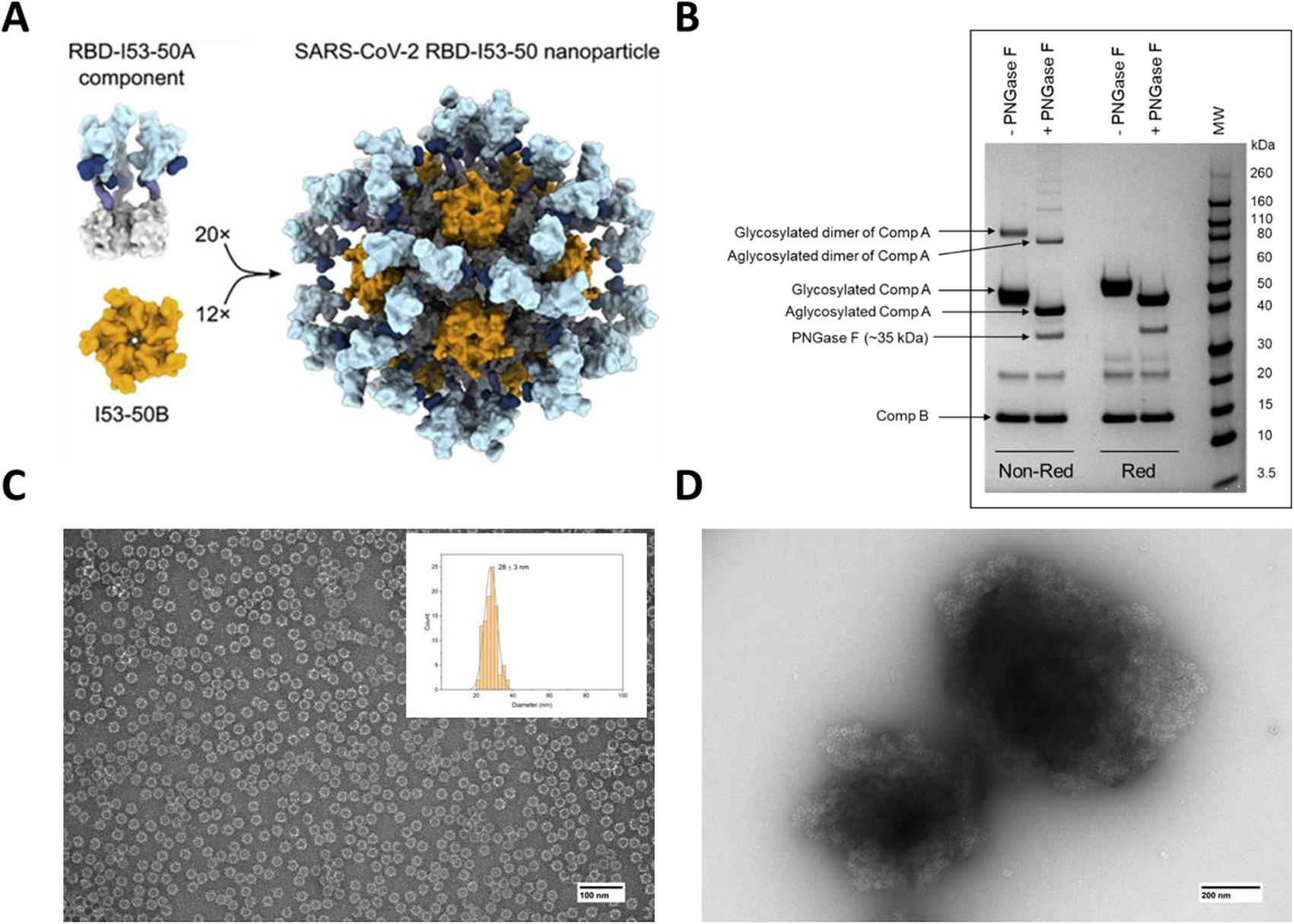
Analysis of IVX-411 VLP. (A) Schematic representation of the two-component IVX-411 VLP (also see text). (B) Representative reducing and non-reducing SDS-PAGE analysis of Components A and B of IVX-411 following treatment with or without PNGase-F. (C) Representative negative stained TEM of IVX-411 in solution with particle size distribution indicated in the insert (n=100 VLPs, mean particle size ± SD = 28 ± 3 nm). (D) TEM analysis of IVX-411 antigen adsorbed to the aluminum-salt adjuvant Alhydrogel. Samples were formulated in a buffer containing 50 mM Tris, 150 mM NaCl, 5% sucrose, 0.1 M L-arginine, pH 8.0. The cartoon display in panel (A) was reproduced from Walls et al ^31^.

Overall secondary and tertiary structure analysis of proteins compromising the IVX-411 antigen was performed by fourier transform infrared (FTIR) and intrinsic fluorescence (IF), respectively. The second-derivative FTIR spectra of the Amide I region (1700-1600 cm^-1^) indicates the secondary structure content is a mixture of α-helix and β turn/sheet consistent with the expected structure of the components of IVX-411 (**Figure 2A**). IF analysis (excitation at 295 nm) showed an emission peak maximum of 332 nm, a result indicating the average tryptophan residue in both components (including RBD) is in a relatively apolar environment (**Figure 2B**). For particle size determination of IVX-411, sedimentation velocity analytical ultracentrifugation (SV-AUC) analysis revealed three different species with the major ∼36S species (∼77%) presumably intact, assembled nanoparticles (**Figure 2C**). Two low abundant species at approximately 9S and 53S were also observed, which were likely impurities and higher ordered species, respectively. An estimated MW of roughly 4.1 MDa was calculated for IVX-411 from the SV-AUC analysis, a result close to the predicted MW of 3.9 MDa based on the primary sequence.

**Figure 2.**
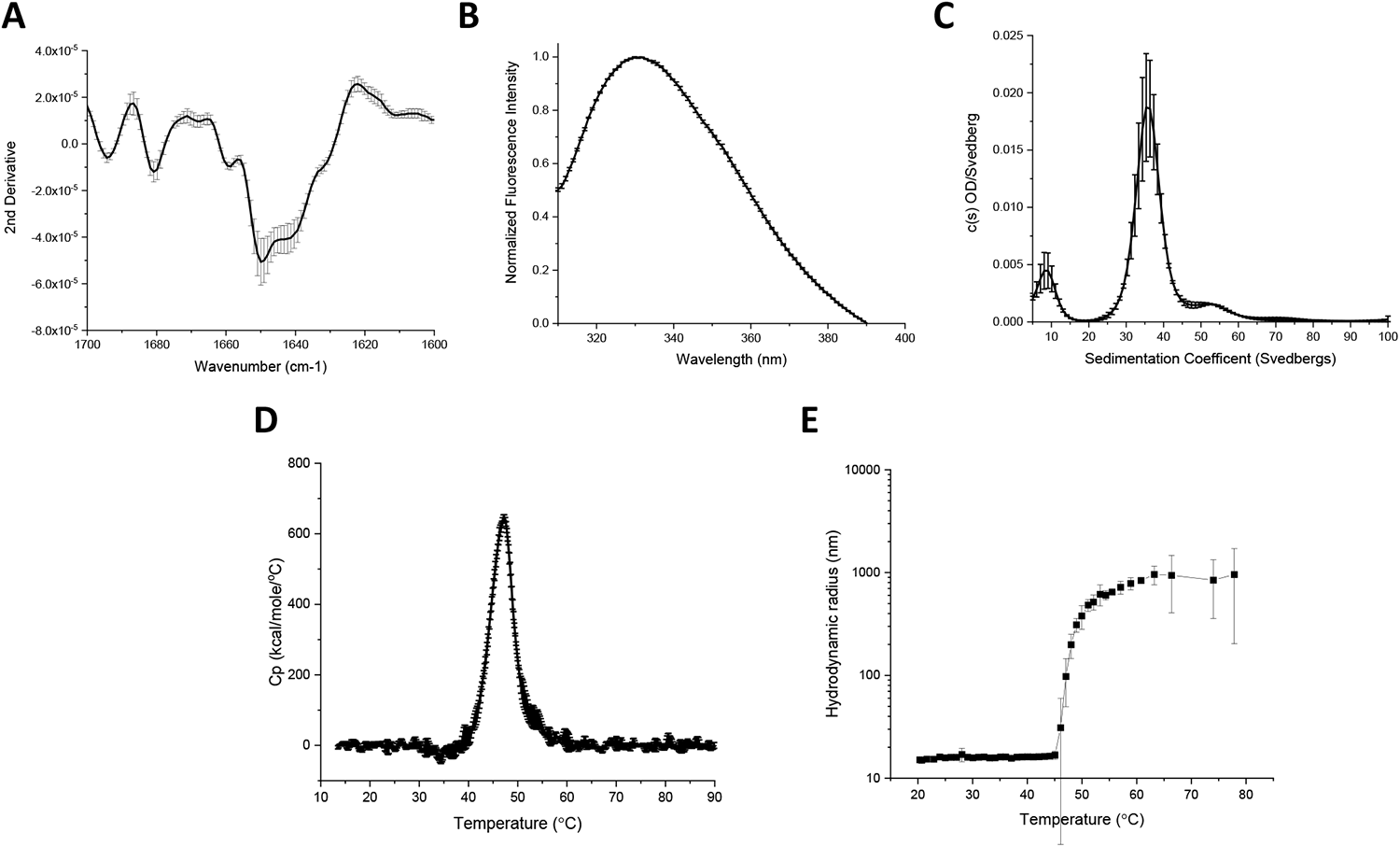
Biophysical characterization of the IVX-411 VLP. (A) Second derivative FTIR spectra of the Amide I band, and (B) Intrinsic Tryptophan emission spectra (excitation at 295 nm) of IVX-411 sample at 10°C. (C) Molecular size distribution of VLP species as measured by SV-AUC. (D, E) The overall conformational stability and aggregation behavior as a function of increasing temperature as measured by DSC (Tm = 47.5 ± 0.1) and DLS (Tm = 48.6 ± 0.7), respectively. Data shown are the mean of at least three independent measurements while the error bars represent one standard deviation. All experiments were performed in a 50 mM Tris, 150 mM NaCl, 5% Sucrose, 0.1 M L-Arginine, pH 8.0 formulation buffer.

The thermal stability of IVX-411 was evaluated by differential scanning calorimetry (DSC) and dyamic light scattering (DLS). A single endothermic event was observed by DSC that initiated at approximately 42°C (onset temperature, Tonset) with a thermal melting temperature of ∼48°C (**Figure 2D**). The aggregation profile of IVX-411 was measured by DLS as a function of temperature (**Figure 2E**). A steep increase in the hydrodynamic radius from ∼14 nm to ∼1000 nm, which is indicative of aggregation, was observed initiating at ∼45°C. Taken together, the DSC and DLS results indicated structural alterations within the IVX-411 antigen initiated at ∼42°C followed by aggregation at ∼45°C.

### IVX-411 antigen binds both AH and AP adjuvants, but with greater binding strength to AH

Prior to evaluating the binding of IVX-411 to AH and AP the point of zero charge was measured by phase analysis light scattering and was ∼5.8 (data not shown). When combined with AH (pI ∼11) in a formulation buffer at pH 8, IVX-411 bound 100% to AH, a result not unexpected if binding occurred via electrostatic interactions. Surprisingly, IVX-411 also bound 100% to AP adjuvant (pI ∼5) in the same formulation buffer (**Figure 3A-B**), suggesting IVX-411 bound to AP via other mechanisms than solely electrostatic. To better understand these observations, unassembled components A and B were evaluated for their ability to bind AH and AP under the same conditions as IVX-411. Both bound nearly 100% to AH, but only bound ∼30% and 56%, respectively, to AP (Figure 3A-B). This result suggests that the binding of IVX-411 to each of the alum adjuvants (AH and AP) is mediated by both components of the mature, two-component VLP.

**Figure 3.**
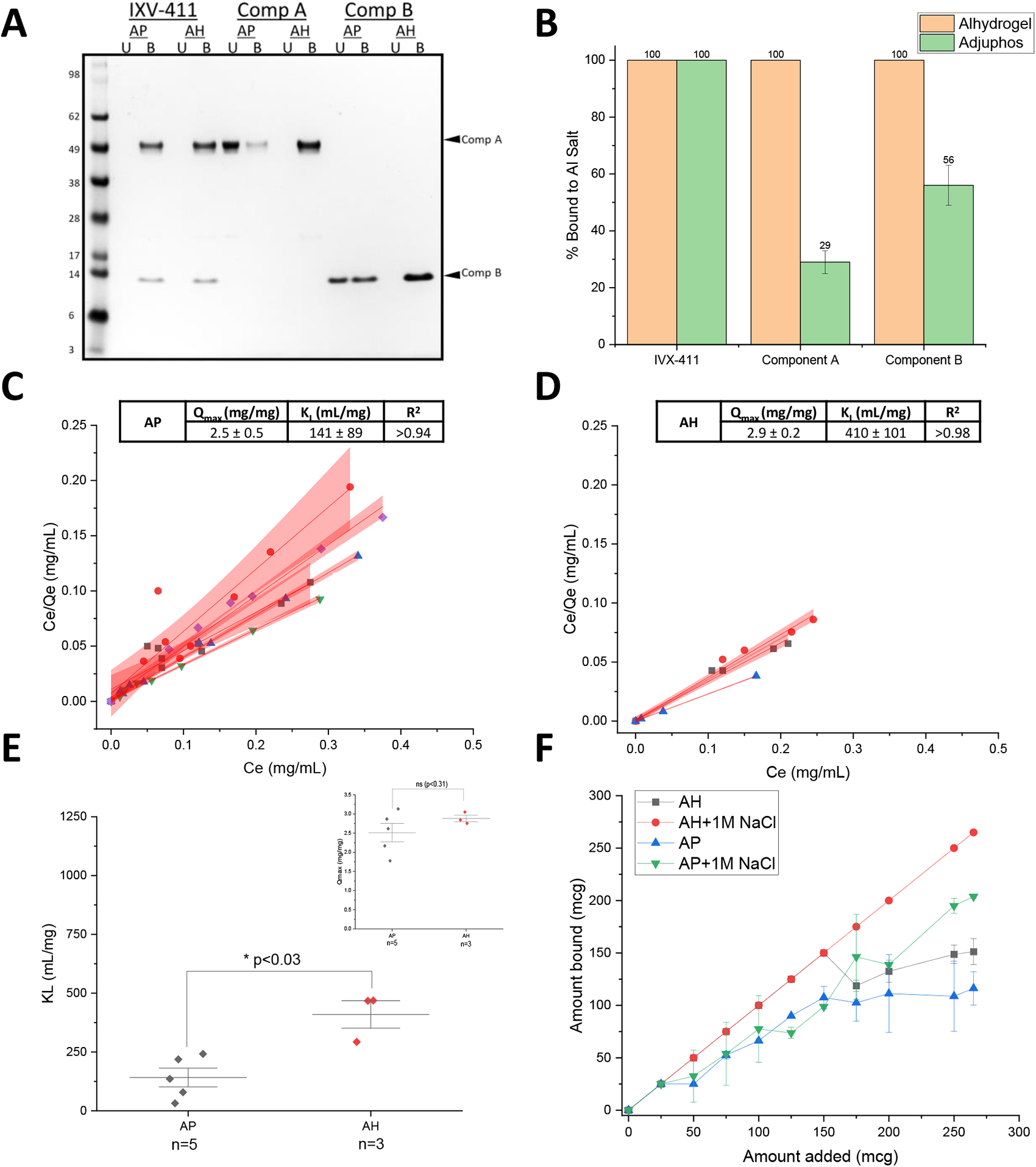
IVX-411 binds to both Alhydrogel (AH) and Adjuphos (AP) while the individual components bind completely to AH and partially to AP. (A) Representative reducing SDS-PAGE of binding of each component and the assembled VLP to AH and AP (U= Unbound; B= Bound). (B) The percentage of each component that is bound to either AH or AP was determined by gel densitometry. The data are the mean of three independent binding experiments with the error bars representing the standard deviation. (C, D) Increasing amounts of IVX-411 (0-600 mcg) were added to a fixed amount (50 mcg) of alum adjuvant (C) Alhydrogel, AH or (D) Adjuphos, AP and the amount of unbound antigen was determined by UV-Visible spectroscopy. The data from individual experiments were each fit using the Langmuir equation (red lines) with the red shading indicating the 95% CIs of the linear fits. (E) Statistical comparisons of the experimentally determined binding strength (K_L_) and binding capacity (Q_max_) values of IVX-411 when bound to AP and AH by a two tailed student’s t-test. The K_L_ and Qmax data are the mean of 3 or 5 binding isotherms for AH and AP, respectively, ± one standard deviation. (F) Addition of 1M NaCl enhances the binding of IXV-411 to AH and AP and could no longer be fit to the Langmuir equation. The data are the mean of two independent experiments with the error bars representing the data range. All experiments were performed in a 50mM Tris, 150mM NaCl, 5% Sucrose, 0.1 M L-Arginine, pH 8.0 formulation buffer.

We determined the monolayer binding capacity and binding strength of IVX-411 to AH and AP via Langmuir binding isotherm studies (**Figure 3C, D)**. The monolayer binding capacity (Qmax) values of IVX-411 to AH and AP were similar, 2.9 and 2.5 mg IVX-411/mg Al salt, respectively, which was statistically insignificant by a two tailed t-test (p<0.3; **Figure 3E inset**). The mean binding strength (K_L_) values, however, differed by approximately three-fold for AH (∼410 mL/g) compared to AP (∼141 mL/g), indicating tighter antigen binding to AH, a result which was statistcically significant by a two-tailed t test (p<0.03) (**Figure 3E**). When the NaCl concentration in the formulation buffer was increased to 1M, more IVX-411 bound to AH and AP, and the binding data could no longer be fit to the Langmuir equation using the same experimental parameters at lower ionic strength (0.15 M) (**Figure 3F**). This latter result indicates that electrostatic interactions are not the primary driver in the binding of IVX-411 antigen to alum (AH or AP) adjuvants, and other molecular interactions are likely primarily responsible for antigen adsorption (see Discussion).

### AH and CpG adjuvants conformationally destabilize IVX-411 while AP and the oil-in water emulsion have minimal effect

Having previously observed conformational destabilization of monomeric RBD when co-formulated with alum and/or CpG adjuvants ^41^, one objective in this work was to determine if multimeric presentation of RBD in a VLP-based antigen had the same destabilizing effects. First, IVX-411 at the protein concentration used in the *in vivo* immunogenicity studies (2 mcg/mL) was formulated with the various adjuvants and antigen-adjuvant interactions were analyzed by SDS-PAGE (**Figure 4A**). As expected, IVX-411 was 100% bound to AH or AP adjuvants independent of the addition of CpG. When IVX-411 was co-formulated with Alum and CpG (5:1 Alum to CpG by weight), approximately 10% and 100% of the CpG was bound to AP and AH, respectively. In the oil-in-water emulsion (AV) formulation, the component A and B bands displayed an apparent slower migration in the gel, likely due to the sample’s higher viscosity (**Figure 4A**.)

**Figure 4.**
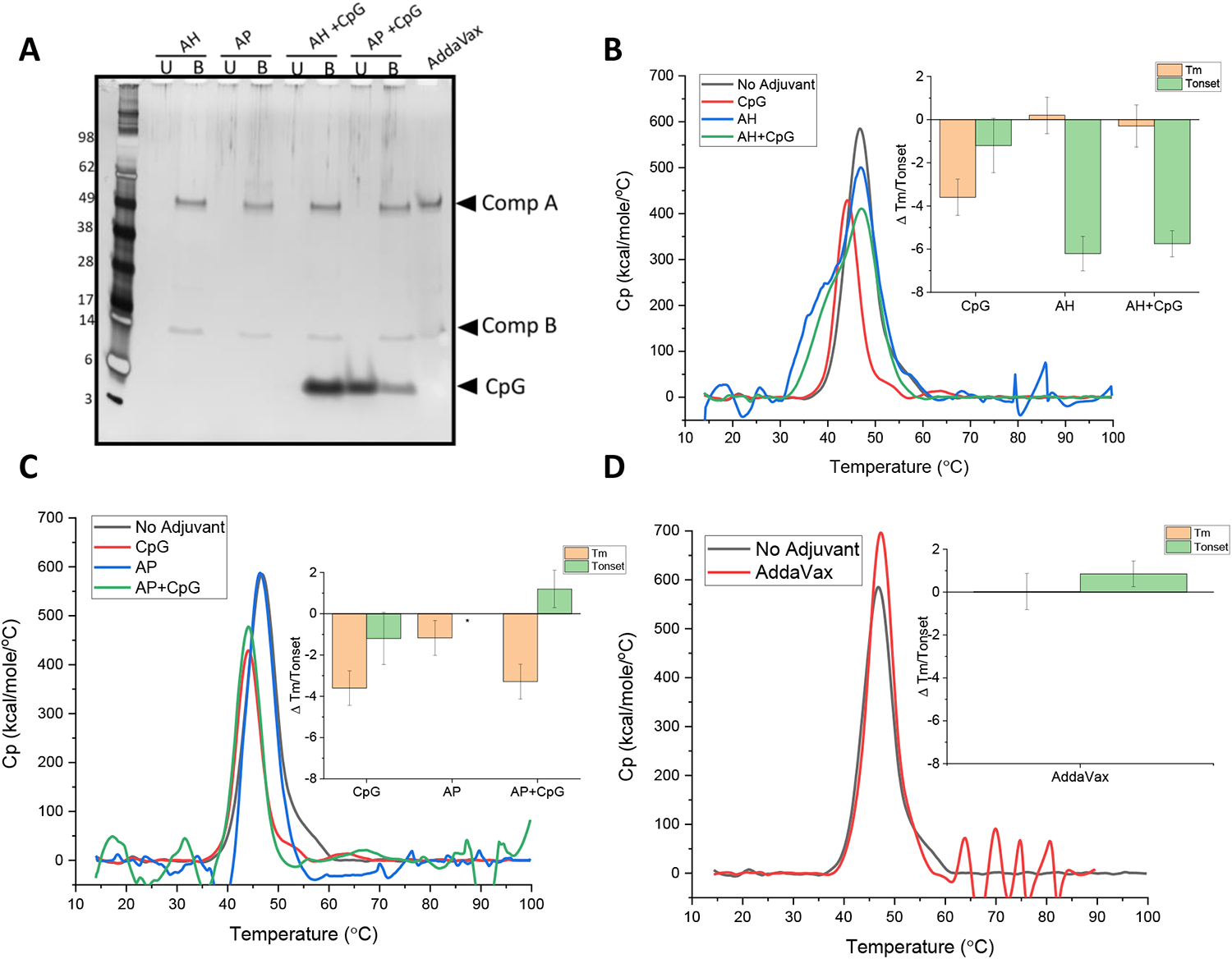
Effect of various adjuvants on the overall conformational stability of IVX-411. (A) Percent antigen bound vs unbound to adjuvant as determined by SDS-PAGE analysis with representative gel showing IVX-411 antigen samples formulated with various adjuvants (2 mcg/mL IVX-411 in presence of AH, AP, with or without CpG, or AV). The unbound **(U)** and the bound **(B)** fractions were separated by centrifugation followed by removal of alum-bound antigen (and CpG) using “strong” desorption conditions (see methods). (B, C) DSC thermograms comparing IVX-411 in solution with IVX-411 formulated with (B) AH or (C) AP alum-adjuvants, both with or without CpG. *the onset temperature value of antigen with AP alone could not be calculated due to baseline noise. (D) DSC thermograms comparing IVX-411 in solution to IVX-411 formulated with AV. As shown in the inset bar graphs, the effect of the adjuvants on the overall conformational stability of IVX-411 was summarized by calculating delta Tm and delta Tonset values (i.e., subtracting the Tm and Tonset values of the no adjuvant control from the Tm and Tonset values of formulated samples). Thermograms are a mean of duplicate scans and the data in the inserts of B-D are the mean of two measurements and the error bars represent the data range after accounting for error propagation. All DSC experiments were performed at 200 mcg/mL IVX-411 in a 50 mM Tris, 150 mM NaCl, 5% Sucrose, 0.1 M L-Arginine, pH 8.0 formulation buffer.

The conformational stability of IVX-411 formulated with the various adjuvants was then assessed by DSC. Compared to unadjuvanted IVX-411, the addition of CpG resulted in a ∼4°C decrease in Tm, without affecting Tonset, values. In contrast, when IVX-411 was formulated with AH, the value of Tonset decreased by ∼6°C with no change in Tm. When IVX-411 was formulated with AP, only a slight change in Tm value was observed (compared to the unadjuvanted control), but Tonset values could not be reliably determined due to baseline noise. When IVX-411 was co-formulated with CpG + AH or CpG +AP, none to small additional destabilizing effects of CpG were observed in the DSC thermograms, respectively (**Figure 4B-C**). Taken together, IVX-411 was 100% bound to both AH and AP, but displayed more structural destabilization when adsorbed to AH (compared to AP) as measured by DSC analysis. Although CpG alone destabilized the overall conformational stability of IVX-411, relatively small, additional destabilizing effects of CpG were observed in the presence of alum. Finally, when formulated in the AV emulsion, no notable changes in Tm or Tonset values of IVX-411 were observed compared to unadjuvanted control (**Figure 4D**).

### Real-time, accelerated and stressed stability studies of adjuvanted IVX-411 formulations as measured by competitive ELISAs

To determine the relative stability of the various adjuvanted IVX-411 formulations (**Table 1**), we developed competitive binding ELISAs with three different ligands: (i) ACE2 receptor, a well-established ligand which binds to neutralizing epitopes on RBD ^42^, (ii) mAb S309, a conformational-dependent, neutralizing mAb that binds adjacent to the ACE2 site ^43^, and (iii) mAb CR3022, a conformational-dependent, non-neutralizing mAb that binds on the opposite side of the ACE2 receptor binding surface ^44^. Briefly, in this assay format, the ligands are preincubated with the formulated IVX-411 samples, centrifuged, and the free ligand concentration (not bound to antigen) is determined by measuring its binding to monomeric RBD absorbed on the ELISA assay plate (see Supplemental methods and as described previously ^41^). Representative binding curves of unadjuvanted IVX-411 and IVX-411 formulated with AH (with and without 37°C incubation for two weeks) with each of the three ligands are displayed in **Figure 5A-F**.

**Table 1:** Summary of the composition of the various adjuvanted IVX-411 formulations used in mouse immunogenicity studies. AH, aluminum hydroxide (Alhydrogel™); AP, aluminum phosphate(Adjuphos™); CpG, CpG-1018™ oligonucleotide, AV: AddaVax™ (oil-in-water emulsion).

| Formulation | IVX-411 antigen dose (mcg) | CpG adjuvant dose (mcg) | Alum adjuvant dose (mcg) | Injection volume (mL) |
| --- | --- | --- | --- | --- |
| AH | 0.2 | 0 | 150 | 0.1 |
| AP | 0.2 | 0 | 150 | 0.1 |
| AH+CpG | 0.2 | 30 | 150 | 0.1 |
| AP+CpG | 0.2 | 30 | 150 | 0.1 |
| AV | 0.2 | 0 | 0 | 0.1 |

**Figure 5.**
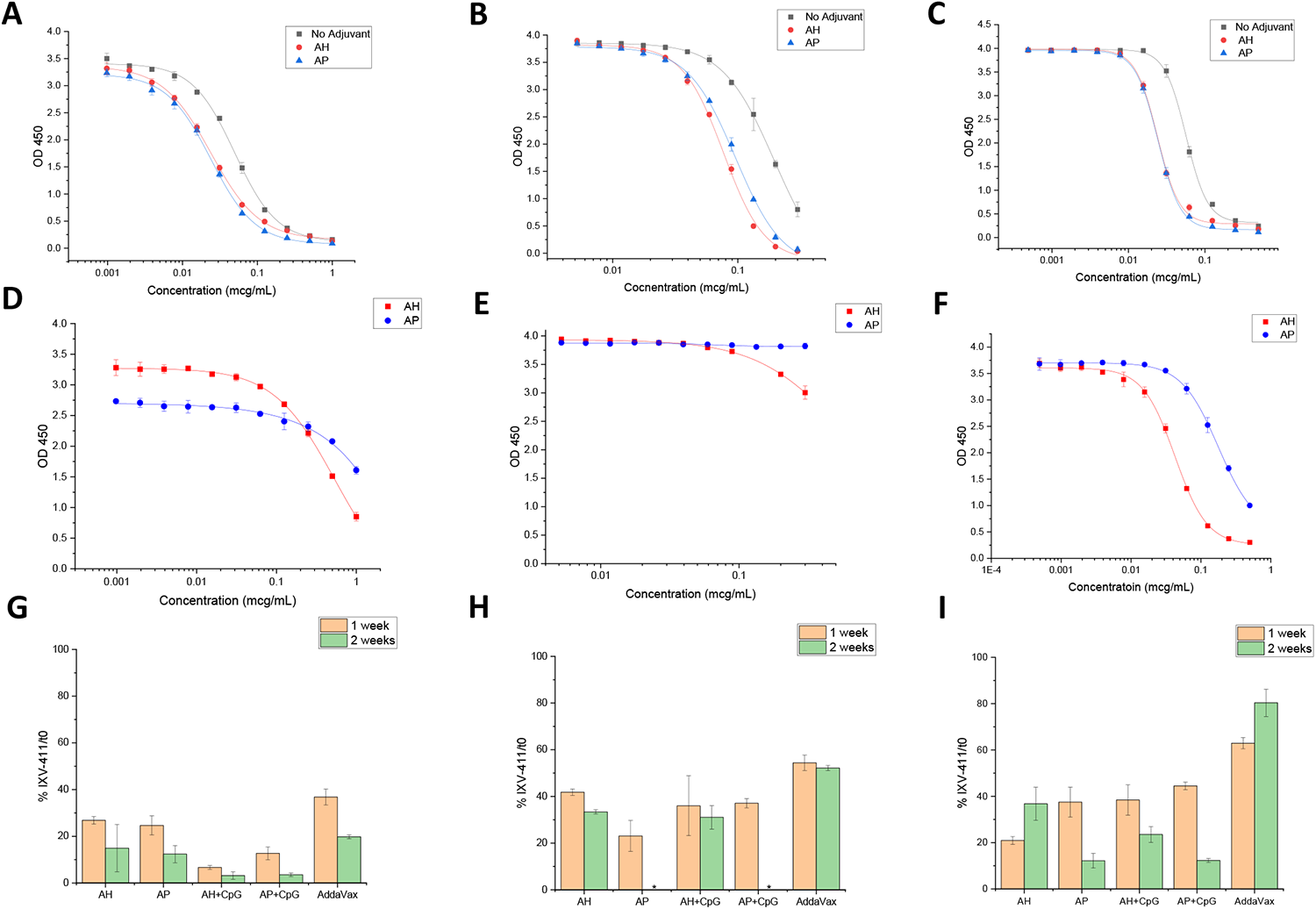
*In vitro* binding assays of IVX-411 in presence of various adjuvants before and after thermal stress. Representative competitive ELISA binding curves of (A) ACE2, (B) S309, and (C) CR3022 ligands to IVX-411 antigen in solution or formulated with either AH or AP. Representative ELISA binding curves of IVX-411 formulated with either AH or AP after incubation at 37°C for two weeks as measured by binding to (D) ACE2, (E) S309, and (F) CR3022. Summary of the relative *in vitro* binding results of formulated IVX-411 samples after incubation at 37°C for one and two weeks for each ligand (G) ACE2, (H) S309, and (I) CR3022. Data shown in G-I are the mean of two samples measured in duplicate (n=4) with the error bars representing one standard deviation. *indicates that no protein was detected. Samples contained 2 mcg/mL IVX-411 formulated with AV, 1.5 mg/mL AH, 1.5 mg/mL AP, 1.5 mg/mL AH + 0.3 mg/mL CpG, or 1.5 mg/mL AP + 0.3 mg/mL CpG.

To further establish the stability-indicating nature of this competitive ELISA assay, the various adjuvanted IVX-411 samples were incubated at 37°C for one to two weeks (**Figure 5G-I**). When compared to 4°C control samples, losses were observed in IVX-411’s ability to bind each of the three ligands after thermal stress. For example, after thermal stress, IVX-411 binding to the ACE2 receptor displayed a ∼80-90% loss vs. binding to the CR3022 and S309 epitopes (60-80% loss). When comparing different formulations, the AV formulation of IVX-411 displayed the slowest degradation rate as measured by the binding to each of the three ligands.

Real-time (2-8°C) and accelerated (25°C) storage stability studies that lasted up to six months were then set up for five IVX-411 adjuvanted formulations (**Table 1**) and the relative stability profiles were monitored by competitive ELISA (**Figure 6**). When compared to ACE2 receptor binding (**Figure 6A-E**), the CR3022 (**Figure 6F-J**) and S309 (**Figure 6K-O**) mAb binding profiles displayed less structural changes in the IVX-411 antigen during longer-term storage at 4 and 25°C, a result consistent with the short-term incubations at 37°C described above. For the alum containing formulations, IVX-411 retained approximately 70-80% of antigen binding relative to time zero after storage at 4°C up to six months as measured by all three competitive ELISAs. At 25°C, loss of antigen binding appeared multiphasic with a more rapid loss during the initial 30-60 days. Overall, the IVX-411 stability profiles at 25°C appeared similar across the alum formulations with ∼20-40% of antigen binding remaining after six months. In contrast, the AV formulation of IVX-411 showed the best stability profile at 4°C and 25°C across the three competitive ELISAs (**Figure 6E, J, O**). For example, the AV-formulated IVX-411 maintained roughly 80-100% antigen binding (relative to time zero) with each ligand during storage at 4°C up to six months. At 25°C, approximately 40% of antigen binding was observed with the ACE2 receptor and approximately 80-90% with the CR3022 or S309 mAb. In summary, the oil-in-water emulsion-based AV formulation of IVX-411 was the most stable during storage at elevated temperatures, and there were no notable differences in the stability profiles of the alum containing formulations independent of co-formulation with CpG.

**Figure 6.**
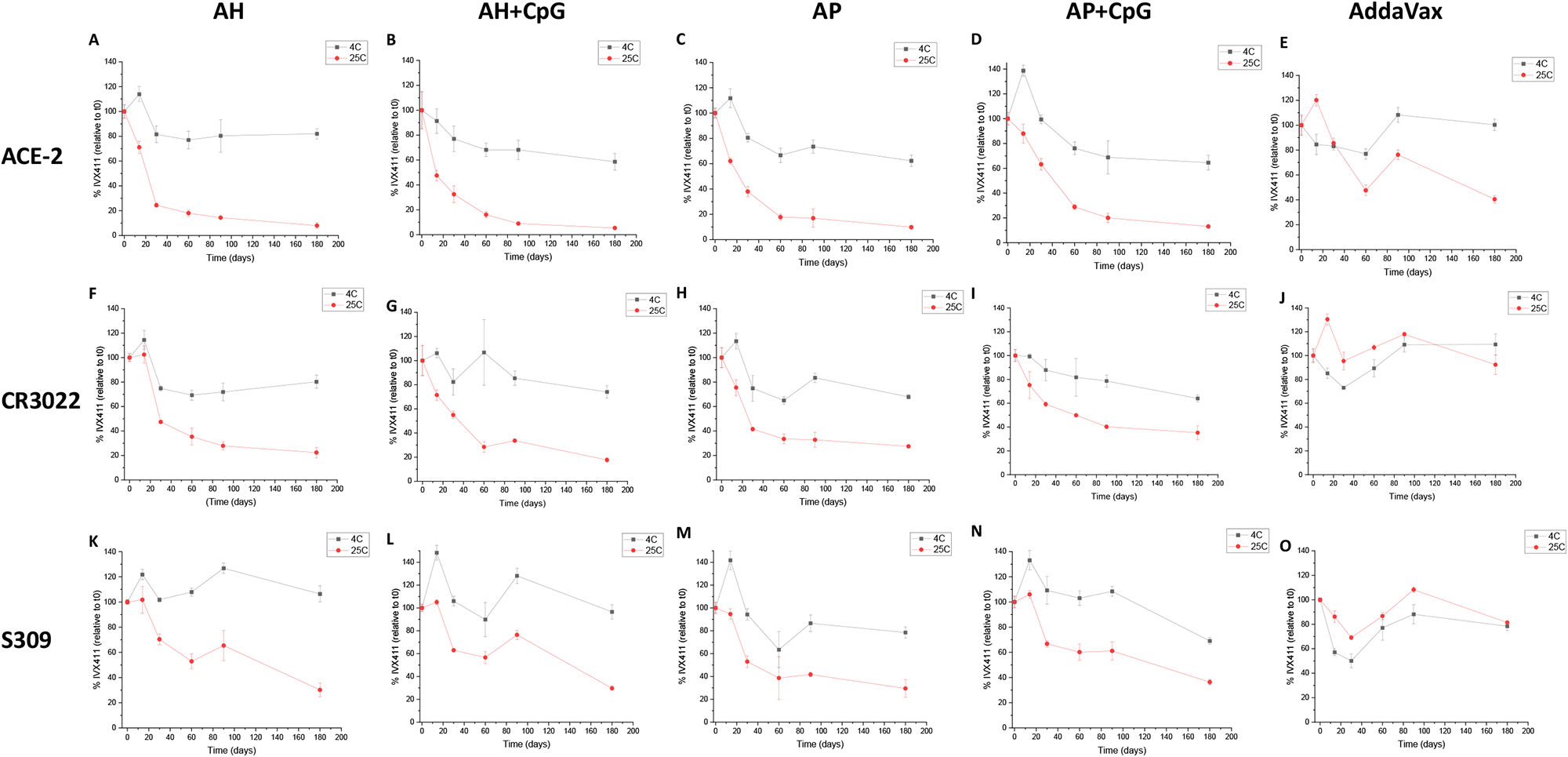
Real-time, accelerated and stressed stability studies of IVX-411 in presence of various adjuvants as measured by *in vitro* binding assays. Competitive ELISA results with formulated IVX-411 samples as measured by binding to (A-E) ACE2, (F-J) CR0322, and (K-O) S309. Samples were stored at 4°C, 25°C, and 37°C and consisted of 2 mcg/mL IVX-411 formulated with either AV, 1.5 mg/mL AH, 1.5 mg/mL AP, 1.5 mg/mL AH + 0.3 mg/mL CpG, or 1.5 mg/mL AP + 0.3 mg/mL CpG. Data shown are the mean of two samples measured in duplicate (n=4) with the error bars representing one standard deviation.

### Effect of IVX-411 dose and adjuvant formulation on mouse pseudovirus neutralization titers

As an initial evaluation of mouse immunogenicity profiles of the adjuvanted IVX-411 formulations, we examined the dose effect of IVX-411 antigen on pseudovirus neutralizing antibody titers (pNT). We immunized mice with 0.2 or 0.02 mcg IVX-411 (total protein) adjuvanted with either 150 mcg AH or AP and co-formulated with 30 mcg CpG. Mice were primed on Day 1 with a booster dose administered on Day 21 (Figure 7A). On Day 21, prior to the booster dose, neutralizing titers at both doses in all formulations were below levels required to generate NT_50_ values (**data not shown**). On Day 35, after the booster dose, we observed NT_50_ values >1 log in the 0.2 mcg dose independent of the formulation (**Figure 7**). This trend continued on Day 65, where the NT_50_ values increased ∼ 1 log further in both formulations and at both doses. No significant differences in NT_50_ values were observed between AH+CpG vs AP+CpG formulations on days 35 or 65 (**Figure 7B**).

**Figure 7.**
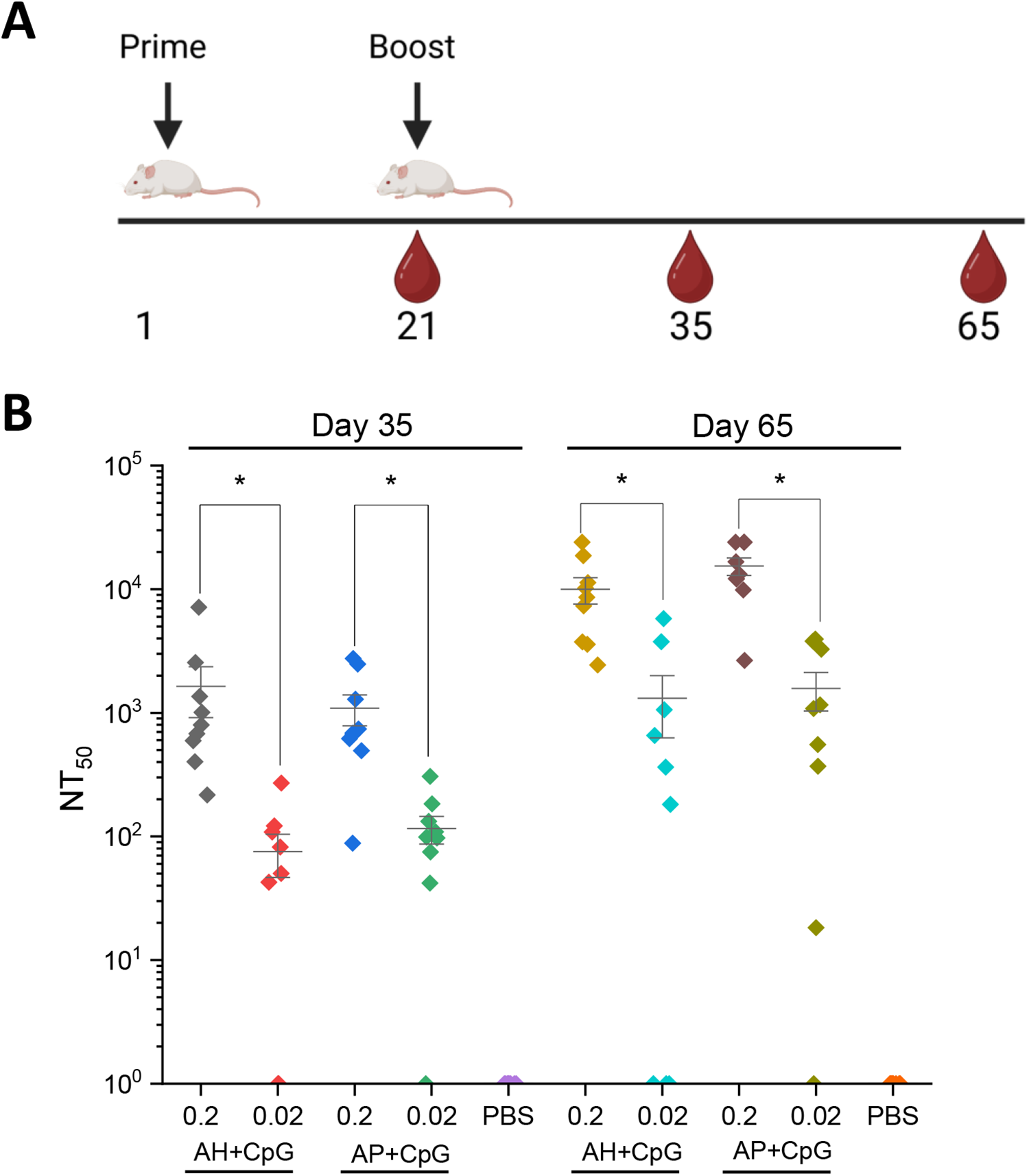
SARS-CoV-2 pseudovirus neutralization activity in the sera of mice immunized with low and high doses of IVX-411 formulated with CpG 1018 and AH or AP. (A) Groups of mice (n=9/group) were immunized on study day 0 with 0.2 or 0.02 mcg IVX-411 adjuvanted with either 150 mcg AH and 30 mcg CpG or 150 mcg AP and 30 mcg CpG. Mice were boosted with a second dose on Day 21. Serum samples were collected on days 35 and 65. (B) NT50 values from sera collected on Days 35 and 65 using SARS-CoV-2 (D614G B.1, 20A) reporter virus particles. Asterisks indicates a statistically significant difference by Kruskal-Wallis analysis using Dunn’s multiple comparisons test (*p<0.03). The diagram in (A) was created using BioRender.

In a second mouse immunogenicity study, various alum adjuvanted formulations of IVX-411 were compared head-to-head with the AV formulation. In addition, to correlate the results of the *in vitro* competitive ELISA results to *in vivo* performance, the IVX-411 formulations were stored at 4°C or 37°C for 2 weeks prior to immunization. The IVX-411 antigen (0.2 mcg dose) was adjuvanted with either AH or AP, both with and without CpG using the same prime-boost dosing regimen described above (**Figure 8A**). Similar to the dose-ranging results described above, NT50 values could not be determined at Day 21 (data not shown.)

**Figure 8.**
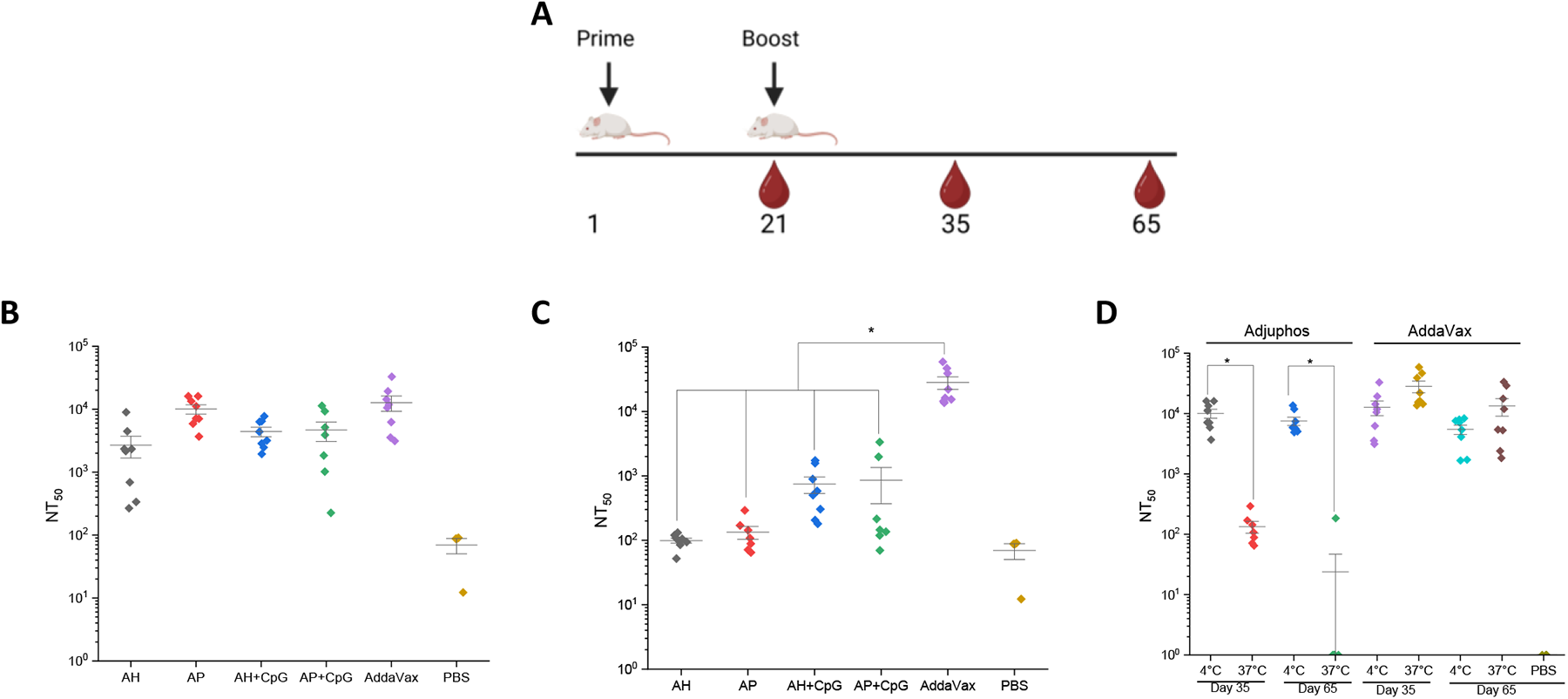
SARS-CoV-2 pseudovirus neutralization titers of mice immunized with IVX-411 in presence of various adjuvants with and without thermal treatment. **(A) Groups of mice** (n=8/group) were immunized with 0.2 mcg IVX-411 along with either 150 mcg AH, 150 mcg AP, 150 mcg AH + 30 mcg CpG, 150 mcg AP+ 30 mcg CpG, or AV. The vaccine formulations were either stored at (B) 4°C or (C) 37°C for 2 weeks prior to immunization and Day 35 neutralization titers were measured. (D) NT50 values derived using SARS-CoV-2 (D614G B.1, 20A) and sera from the AP and AV formulations at Days 35 and 65 were compared after storage at 4°C and 37°C for 2 weeks. Mice were boosted on Day 21 and sera collected on Days 35 and Asterisks indicates a statistically significant difference by Kruskal-Wallis analysis using Dunn’s multiple comparions test (*p<0.03). The diagram in (A) was created using BioRender.

On Day 35, the highest neutralizing titers were observed in the IVX-411 groups formulated with either AV or AP (**Figure 8B**). Approximately 2-4 fold lower neutralization titers were observed when the AV group was compared to the other groups (AH, AH+CpG and AP+CpG). For the IVX-411 formulations stored at 37°C for 2 weeks, at least a 5-fold decreases in neutralization titers were observed in all formulations except for the AV-adjuvanted formulation, which displayed a trend of slightly higher titers relative to the 4°C samples (**Figure 8C**). When comparing the alum adjuvanted groups of IVX-411, with or without CpG, no differences in pnt values were observed with non-stressed samples. After thermal stress, although a ∼6-7 fold higher mean neutralization titers were observed in the alum + CpG containing formulations (vs alum alone), these differences were statistically insignificant (p>0.9, a result due to the two mice in each group that generated pNT >1000, which influenced the calculated the mean values).

We further compared the neutralizing titer responses of the two best performing IVX-411 formulations, adjuvanted with either AP or AV, at day 35 and day 65. The levels of neutralizing antibodies were sustained through day 65 in both formulations (**Figure 8D**). When incubated at 37°C for two weeks prior to immunization, there was, however, a ∼500 fold decrease in the neutralization titers generated by the AP (versus AV) adjuvanted formulations (Figure 8D). In summary, for the IVX-411 adjuvanted formulations stored at 4°C, the AV or AP formulations displayed statistically similar, and the highest levels of, neutralizing titers in mice. The AV-formulated IVX-411, however, maintained the ability to induce a relatively high neutralizing antibody response after storage at 37°C for two weeks whereas the capacity of the alum adjuvanted formulations to elicit neutralizing antibodies were greatly diminished.

## Discussion

The major goal of this work was to assess the feasibility of developing a second-generation, low-cost COVID-19 vaccine candidate containing an aluminum-salt adjuvanted formulation of a two-component VLP displaying SARS-CoV-2 antigen. To this end, we optimized aluminum-salt (i.e., AH, AP) adjuvanted formulations of IVX-411, and then compared their stability profiles and *in vivo* performance to IVX-411 formulated with newer adjuvant (i.e., alum+ CpG, oil-in-water emulsion, AV). In addition, results from the RBD-antigen displayed within a VLP (IVX-411) from this work can be compared to our recently reported results with a monomeric RBD antigen alone (RBD-J) formulated with the same alum adjuvants with and without CpG ^41, 45^.

### Optimizing aluminum-salt formulations of IVX-411

Differences in antigen-aluminum adjuvant binding strength can play a key role in immunopotentiation. If the adsorptive strength is too tight, it could potentially cause interfere with antigen processing and presentation to antigen presenting cells via MHC-II and thus impede a robust B-cell response. We observed reduced pNT values in mice immunized with IVX-411 adjuvanted with AH alone when compared to AP alone. Interestingly, the IVX-411 antigen in the formulation buffer completely bound to both AH and AP aluminum-salt adjuvants. When the binding interactions were quantified by Langmuir binding isotherm analysis, there was approximately 3-fold greater binding strength of IVX-411 to AH compared to AP. This result suggests increased binding strength of IVX-411 to AH may have resulted in reduced immunopotentiation. Hansen et al. observed similar results with decreased antibody responses of a vaccine antigen as a function of increasing adsorption strength to alum ^46^. Similarly, increased immunogenicity was observed when Hepatitis B surface antigen (HBsAg) was less strongly bound to alum adjuvant ^47^.

In the formulation buffer at pH 8, the surface of the colloidal suspension of AH is positively charged while AP is negatively charged ^13^. IVX-411 is composed of two components that have different isoelectric points, Comp A and Comp B with pI values of 8.7 and ∼6.7, respectively, as calculated from primary structure. Thus, it is possible that components A and B could be binding AP and AH, respectively, through preferential electrostatic interactions at pH 8. When we evaluated these individual components, however, we observed complete binding to AH. Binding of both individual components to AP, albeit incomplete, was also observed. Thus, it is unlikely that IVX-411 binds to AH and AP via a bipolar and strictly electrostatic mechanism as reported by Dagouassat et al ^48^ for BBG2Na, which is another multicomponent vaccine (see below). On the other hand, it is possible that the increased avidity of IVX-411 through multivalent presentation of RBD could be responsible for binding to both AH and AP.

When the ionic strength of the formulation buffer was increased approximately six-fold (i.e., 0.15 to 1M NaCl), not only did more IVX-411 antigen bound to AH or AP, but the binding results could no longer be fit to a Langmuir model. These observations further suggest that electrostatic interactions are not the primary forces responsible for IVX-411 binding to AH and AP. Higher NaCl concentrations could promote hydrophobic interactions between antigen-adjuvant by reducing solute solvation, increasing exposure of hydrophobic regions of the RBD-VLP, and thus increasing the amount of IVX-411 bound to the colloidal surface of the aluminum-salts (AH, AP) independent of the surface charge ^49, 50^.

We also observed a lowering of the conformational stability of IVX-411 when bound to AH, but not AP, alum adjuvant as measured by DSC. This result is consistent with stronger interactions between IVX-411 and the surface of AH. Although varying degrees of conformational destabilization of different protein antigens after adsorption to aluminum-salt adjuvants have been widely reported ^41, 51–54^, there are exceptions to this trend with other reports observing no notable structural destabilization after antigen adsorption ^55, 56^. Taken together, the antigen-adjuvant binding data suggest hydrophobic interactions are the main driving force responsible for IVX-411 antigen adsorption to aluminum-salt adjuvants, although electrostatic interaction may also play a secondary role that could explain stronger IVX-411 antigen-alum adjuvant interactions with AH compared to AP.

These results, although intriguing, are not unprecedented and have been observed with other vaccine antigens. For example, the fusion protein BBG2Na comprised of the central domain of RSV G glycoprotein, G2Na, and the albumin binding domain of *Streptococcus* protein G, BB, bound both AH and AP ^48^. The mechanism of this binding, however, was primarily electrostatic via the acidic pI of BB (5.5) and the basic pI of G2Na (10.0) that was responsible for binding AH and AP, respectively ^48^. Interestingly, robust *in vivo* immunogenicity and protection profiles in mice were observed with a subunit RSV fusion protein (BBG2Na) vaccine candidate upon RSV challenge independent of the type of aluminum-salt adjuvant used ^48^. Another study found high binding (>90%) of recombinant botulinum toxin fragments to both AH and AP, although the results were solution pH and ionic strength dependent, which the authors proposed were a combination of electrostatic and ligand exchange interactions ^57^. Finally, HPV 16 VLPs have been observed to bind both AH and AP adjuvants ^58^.

### Stability profiles of Alum adjuvanted formulations of IVX-411 (RBD-VLP) compared to other adjuvants and monomeric RBD antigen

The antigen-adjuvant interactions, storage stability profiles, and mouse immunogenicity results described above for alum formulations of IVX-411 were then compared with IVX-411 formulations containing either CpG, alum + CpG (i.e., AH+CpG, AP+CpG) or an oil-in-water emulsion (AV). In addition, we compared these results to our recently reported results examining the stability and immunogenicity of a monomeric RBD antigen (RBD-J) formulated with the same adjuvants ^41^. Originally, we had also planned to compare the stability of IVX-411 in solution to that of RBD-J in solution. This analysis, however, was hindered due to adsorption of IVX-411 to the surface of the glass vial at the low protein doses examined in this work (data not shown). Primary container surface adsorption was not observed when IVX-411 was pre-adsorbed to alum, or formulated with AV adjuvants.

The molecular mechanisms by which adjuvants either stabilize, destabilize, or have no effect on the structural integrity of subunit vaccine antigens remain poorly understood ^51, 52^. Several studies have examined the effects of thermal and freeze induced degradation of protein antigens on the surface of aluminum salts ^25, 52, 53, 59^. In a recent study by our group, we observed that a monomeric RBD-J antigen adjuvanted with AH or AH+CpG (CpG ™ Adjuvant) displayed a dramatic decrease (8-27°C) in conformational stability as measured by DSC ^41^. A roughly 12°C decrease in the Tm value was measured in the previous work when monomeric RBD (RBD-J) was adsorbed to AH, while no notable change in Tm values were observed in this work when multimeric RBD on a nanoparticle (IVX-411) was adsorbed to AH. Moreover, DSC results for IVX-411 upon the addition of CpG alone resulted in an approximately ∼4°C decrease in Tm values, which was a notably less destabilization effect than was observed previously with monomeric RBD-J (∼13°C decrease) ^41^. The RBD-J antigen is a stabilized form of RBD as a result of two amino acid substitutions, so the RBD portion of IVX-411 could be further optimized for greater stability ^27^. These results are consistent with the RBD antigen being more stable when displayed on the surface of a VLP (IVX-411), compared to a monomeric protein (RBD-J), likely due to the nanoparticle component protecting the RBD antigen from direct destabilization via alum binding. Finally, no notable conformational destabilization of IVX-411 was observed in the AV formulation by DSC. This result indicates that the oil-in-water emulsion adjuvant (AV) was the least destabilizing for IVX-411 amoungst the adjuvants analyzed in this study.

We also monitored the storage stability of the various IVX-411 adjuvanted formulations during real-time and accelerated temperature conditions. To this end, we developed an *in vitro* competitive ELISA to monitor the ability of IVX-411 to bind ACE2 receptor and the mAbs CR3022 and S309 as a function of time at three different temperatures with the objective to determine if the adjuvants affected the stability of IVX-411. One advantage of using competitive ELISA is its ability to monitor critical epitopes on RBD that are directly related to antigenicity of the vaccine candidate as opposed to measuring overall conformational changes by DSC. In addition, there is no need to desorb/extract the antigen from alum or the oil-in-water emulsion adjuvants, respectively, which allows for direct measurement of the antigen integrity in the presence of adjuvants.

Over a six-month timeframe during storage at 4° or 25°C, we observed similar IVX-411 stability profiles in the AH, AH+CpG, AP, or AP+CpG formulations. The AV adjuvanted formulation displayed superior stability profile over six months with 80-100% of CR3022 and S309 ligand binding ability remaining after 6 months at 4°C and 25°C. At 37°C, similar losses in antibody binding were observed in all formulations with the AV formulation displaying the best relative stability profile. A similar degradation pattern was observed while studying monomeric RBD (RBD-J) formulated with AH at 4°C and 25°C over the course of 90 days ^41^, but the loss of the ability to bind ACE2 was accelerated when CpG was co-formulated with AH. Conversely, IVX-411 adjuvanted with CpG had no effect on ACE2 binding as a function of time or temperature when co-formulated with alum adjuvants. Taken together, these data suggests that the stability profile of the formulated RBD antigen is stabilized by multimeric incorporation within a VLP (IVX-411) compared to the monomeric form (RBD-J).

### Mouse immunogenicity profiles of aluminum-salt formulations of IVX-411 (RBD-VLP) compared to other adjuvants and monomeric RBD-antigen

Comparable pNT values were measured in mice immunized with AP and AV, while lower titers were observed when IVX-411 was adjuvanted with AH, independent of co-formulation with CpG. These observations are in stark contrast to monomeric RBD-based subunit vaccine candidate (RBD-J), which had a higher neutralizing antibody response when AH was co-adjuvanted with CpG, yet was destabilized during storage ^41, 60^. In the case of IVX-411, the addition of CpG to alum adjuvanted IVX-411 did not improve neutralization titers in mice when vaccinated with non-stressed samples (stored at 4°C). On the other hand, in mice administered stressed IVX-411 samples (37°C, 2 weeks), the same CpG + alum samples showed a trend towards improved neutralization titers (albeit statistically insignificant due high variability). Finally, alum formulations with and without CpG displayed similar IVX-411 stability profiles during real time and accelerated storage.

The IVX-411 doses used in this mouse immunogenicity study were >1 order of magnitude lower (0.2 mcg total antigen which is ∼40% RBD by weight) when compared to monomeric RBD-J formulations used in previous mice studies (5 mcg), a result which indicates IVX-411 induced a more potent immune response than monomeric RBD-antigens. Similar results have been reported with engineered RBD nanoparticles constructed using a SpyTagged RBD and I301 nanoparticles containing SpyCatcher ^27^. Moreover, more robust neutralizing and cell-mediated responses were observed in a RBD/N subunit protein vaccine candidate adjuvated with AH+CpG (ODN2395) when compared to other adjuvant combinations (i.e., MPLA/Quil A, AddaS03) ^61^.

After thermal stress, the loss of IVX-411 binding to ACE2 receptor in the competitive ELISA assay correlated well with the reduction in pNT in mice with alum adjuvanted IVX-411 formulations. For thermally stressed AV-adjuvanted IVX-411 formulations, no reduction in neutralization titers in mice were observed, which correlated with the superior stability profiles at 4°C, 25°C, and 37°C observed in the competitive ELISA using ACE2, CR3022, and S309 ligands. The surfactants polysorbate 80 and sodium trioleate could be responsible for this enhanced stability in the AV formulation ^62, 63^. Excipient optimization was outside the scope of the current study; however, future work could better optimize the AP formulation to enhance IVX-411 storage stability to better maintain vaccine potency through manufacturing, storage, and distribution.

Losses in an *in vitro* potency assay may or may not correlate with *in vivo* immunogenicity. For example, losses in D-antigenicity in an inactivated Sabin poliovirus vaccine subjected to thermal treatment did not correlate with reduced immunogenicity in rats ^64^. Another example includes AH+CpG adjuvanted RBD-J vaccine candidate, which lost roughly 90% of the *in vitro* activity by ACE2 competitive ELISA, but retained the ability to generate robust pNT in mice (after two doses, but to a lesser extent after one dose) ^41^. The same study, however, demonstrated extensive denaturation of the RBD protein (pre-treatment with the reducing agent DTT) resulted in a complete loss of RBD ACE2 binding by competitive ELISA that correlated with the near complete abrogation of a neutralizing antibody response in mice ^41^. Taken together, these results demonstrate that near complete loss of ACE2, CR3022, and S309 binding in the *in vitro* competitive ELISA assay can correlate well with the reduced ability to generate a robust *in vivo* neutralizing antibody response in mice.

## Conclusions and Future Work

One major goal of this study was to optimize a low-cost alum adjuvanted formulation for IVX-411 and compare the results to other widely-used adjuvants. Interestingly, although IVX-411 bound different aluminum-salt adjuvants (i.e., AP and AH) with similar antigen-adjuvant binding capacities, the AP-adjuvanted IVX-411 displayed superior neutralizing titers and weaker antigen-adjuvant binding strength. The AP-adjuvanted IVX-411 was compared to alum+CpG and oil-in-water emulsion (AddaVax, AV) formulations in terms of SARS-CoV-2 pseudovirus neutralizing antibody response in mice. The highest neutralization titers were generated in mice immunized with IVX-411 adjuvanted with either AP or AV. CpG addition to alum adjuvanted IVX-411 had no effect on pNT values when vaccinating with non-stressed samples, but displayed a trend (yet statistically insignificant) of improved mean pNT values in mice immunized with stressed samples stored at 37°C for two weeks. This result differs from our recent report with monomeric RBD antigen (RBD-J) formulated with the same adjuvants, where AH+CpG adjuvanted RBD-J generated the highest pseudovirus neutralization titers, independent of thermal treatment ^41^.

We also compared these adjuvanted formulations of IVX-411 in terms of their relative stability profile as a function of time and temperature by developing competitive ELISAs that monitored antigen binding to the ACE2-receptor (or conformational mAb) in the presence of adjuvants. The AP and AV formulations of IVX-411 displayed similar storage stability profiles at 4°C as measured by competitive ELISAs. Interestingly, IVX-411 adsorption to alum or co-formulation with CpG did not affect storage stability, a result in contrast to our previous report with monomeric RBD which was destabilized by these adjuvants ^41^. These results suggest the adjuvanted formulations of the RBD are more stable when the RBD antigen is within in a multimeric display on the surface of a VLP compared to the monomeric form by itself. It will be of interest in future work to evaluate if other formulated protein antigens display similar differences in their stability profiles in the monomeric form vs when expressed in a multimeric display on the surface of a I53-50-based VLP.

## Abbreviations

AH: Alhydrogel™
AP: Adjuphos™
CpG: CpG 1018™ Oligonucleotide Adjuvant
AV: AddaVax™
pNT: pseudovirus neutralizing titers
TLR: toll-like receptors
RBD: receptor binding domain
GMP: good manufacturing practices mDa megadalton
AS03: Adjuvant System 03
AS04: Adjuvant System 04
AS37: Adjuvant System 37
VLP: Virus-Like Particle
IVX-411: VLP vaccine containing RBD antigen using the I53-50 two component recombinant protein platform

## Acknowledgements

We acknowledge the Bill and Melinda Gates Foundation for funding this work (# INV-021035 and INV-027417.) We thank the Global Health Discovery Collaboratory for supplying the recombinant ACE2-Fc receptor, CR3022 mAb, and the SARS-CoV-2 receptor binding domain used as immunoassay reagents. We thank Dynavax Technologies for providing the CpG 1018™ adjuvant and Prem Thapa in the University of Kansas Microscopy and Analytical Imaging Core Lab for assistance with collecting and processing the TEM images. We also thank Scot Shepard, Ross Taylor, and Prathima Acharya at Icosavax, for thoughtful discussions and reviewing the manuscript.

## Data availability statement

All data associated with this study are available with the corresponding author.

