## Supplemental Methods for "Effects of aluminum-salt, CpG and emulsion adjuvants on the stability and immunogenicity of a virus-like particle displaying the SARS-CoV-2 receptor-binding domain (RBD)"

**Current affiliations:** SB: Merck & Co., West Point, PA, USA

KK: Merck & Co., West Point, PA, USA

GVS: Regeneron, Albany, NY, USA

HK: SK Bioscience, Cambridge, MA, USA

CR: Pharmorosconsulting, Bozeman, Mt USA

**Running title:** Stability and immunogenicity of an RBD-VLP vaccine candidate in different adjuvant formulations

**Keywords:** Vaccine, Formulation, Stability, Aluminum-salt, CpG, Emulsion, Adjuvant, SARS-CoV-2, Antigen, VLP (Virus-Like Particle)

**Dynamic Light Scattering-** For isothermal measurements, IVX-411 was diluted to 0.1 mg/mL in 50mM Tris, 150mM NaCl, 0.1 M L-Arginine, 5% Sucrose, pH 8.0 and centrifuged at 12,000 x g for 5 minutes. The supernatant was removed and 20uL was loaded into a 384 well plate (Corning, Corning, NY). The plate was then centrifuged briefly before loading in a Wyatt DynaPro plate reader III (Wyatt Technologies, Santa Barbara, CA) set to 25°C. For thermal ramping measurements, IVX-411 was diluted to 0.2 mg/mL, centrifuged briefly as above, and sealed with plate sealing tape, placed in the instrument, and the temperature was ramped from 20-80°C using a 1°C/min ramp rate with the laser attenuation set to automatic. The data was analyzed by both cumulants or regularization and corrected for buffer viscosity and temperature using Dynamics v7.10.1.21 software.

**Transmission Electron Microscopy (TEM)-** Five µL of sample containing 0.4 mg/mL IVX-411 and 0.2 mg/mL AH or AP in ultrapure (18.2 MΩ) water was mixed well and adsorbed on a 300-mesh carbon-coated copper grid glow discharged for 30 sec using EMS150R S sputter coater (Electron Microscopy Sciences, USA). After 1 min, the grid was washed with a drop of distilled water and negatively stained with 2% Uranyl Acetate (Sigma) solution for 3 min. The excess solution was wicked out with a Kim wipe and the negatively stained grid was examined using a 200kV Hitachi H8100 thermionic field emission transmission electron microscope at an electron acceleration voltage of 200 kV. TEM images were captured using a normative and standardized electron dose on eucentric specimen stage and a constant defocus value from the carbon-coated surfaces. Images were randomly acquired at 10 different locations within the grid.

**Sedimentation Velocity Analytical Ultracentrifugation-** Sedimentation velocity analytical ultracentrifugation (SV-AUC) experiments were performed using an Optima analytical ultracentrifuge equipped with a scanning ultraviolet-visible optical system (Beckman Coulter, Indianapolis, IN.) All experiments were conducted at 20°C after ≥1 hr of equilibration after the rotor reached the set temperature. A rotor speed of 7,000 RPM, UV detection at 280 nm, and a scan frequency of 60 sec were used. IVX-411 samples at 0.3 mg/ml were diluted in 50mM Tris, 150mM NaCl, 0.1 M L-arginine, 5% Sucrose, pH 8.0 and were loaded into Beckman charcoal-epon two sector cells with 12 mm centerpieces and either sapphire or quartz windows and ultracentrifugation was performed overnight. Formulation buffer alone was loaded in the reference cell. The data were analyzed using Sedfit (Peter Schuck, NIH). The formulation buffer density and viscosity used in the analysis was 1.0305 g/mL and 1.24 centipoise, respectively, and were calculated using Sednterp (Thomas Laue, BITC) based on the formulation buffer composition. A continuous c(s) model was used with 900 scans with every 9$^{th}$ scan loaded in the analysis for a total of 100 scans with a range of 0 to 200 svedbergs. A resolution of 300 points per distribution and a confidence level of 0.95 were applied to all fits. Baseline, radial independent noise, and time independent noise were fit, while the meniscus and bottom positions were set manually. The c(s) distributions were imported into Origin 2020 (OriginLab, Northampton, MA) for peak integrations and graph generation.

**FTIR-** FTIR spectroscopy of IVX-411 at 0.68 mg/mL was performed using a Bruker Tensor-27 FTIR spectrometer (Bruker Optics, Billerica, MA) equipped Bio-ATR sample cell and a KBr beamsplitter. The MCT detector was cooled with liquid $N_2$ for at least 20 min prior to use and the interferometer was constantly purged with $N_2$. Two hundred fifty-six scans were recorded from 4000 to 600 $cm^{-1}$ with a 4 $cm^{-1}$ resolution. Background measurements were acquired with formulation buffer (50mM Tris, 150mM NaCl, 0.1 M L-arginine, 5% Sucrose, pH 8.0) alone and subtracted from the sample spectra. Atmospheric and baseline corrections were applied using OPUS V 8.0 (Bruker Optics, Billerica, MA) software. Calculation of the second derivative of the Amide I band (C=O stretching) was also completed using OPUS V 8.0 while applying a 9-point Savitzky-Golay smoothing function.

**SDS-PAGE-**Protein samples were diluted in LDS sample buffer in the presence or absence of 50 mM DTT (Bio-Rad) for obtaining reduced or non-reduced samples, respectively. The diluted samples were heated at 95˚C for 10 min. Samples were loaded on 4-12% Bis-Tris gel (Thermo-Fisher, Waltham, MA) in MES-SDS running buffer (Thermo-Fisher). Gels were stained with Coomassie Blue (Thermo-Fisher) for at least 1 hr and destained with water. Gels that contained CpG were stained using a Quest Silver Stain kit (Thermo-Fisher) where the fixation step was no longer than 30 min. Gels were imaged using a FluorChem E imaging system (Bio-Techne, Minneapolis, MN). For deglycosylation studies, the protein was treated with PNGase F (NEB, Ipswich, MA) per the manufacturer's protocol prior to SDS-PAGE.

**Intrinsic Fluorescence spectroscopy -**The intrinsic tryptophan fluorescence of IVX-411 was measured in triplicate using a dual emission PTI QM-40 Spectrofluorometer (Horiba, Birmingham, NJ) equipped with a 4-position cell holder peltier temperature control device. Fluorescence emission spectra of 0.1 mg/mL IVX-411 from 310-390 nm using an excitation wavelength of 295 nm (>95% Trp) with a step size of 1 nm and an integration time of 1 sec. The excitation and emission slits were both set to 3 nm. All spectra were corrected for background buffer fluorescence and were collected using FelixGX software (Horiba).

**Differential Scanning Calorimetry-** IVX-411 samples were diluted to 0.2 mg/mL in 50mM Tris, 150mM NaCl, 0.1 M L-arginine, 5% Sucrose, pH 8.0 and loaded in a DSC autosampler tray held at 4ºC. DSC was performed using an Auto-VP capillary differential scanning calorimeter (Malvern, Northhampton, MA) using pressurized cells at ~65 psi.  Two water-water scans were taken prior to the first reference (buffer-buffer) scan used for reference correction. Scans were completed from 10-100ºC using a scan rate of 60ºC/h, a pre-scan thermostat of 15 min, and a filtering period of 10. Reference subtraction and concentration normalization were performed using the instrument software.

**Competitive ELISA-** Samples were diluted to 1 mcg/mL, 0.5 mcg/mL, or 0.3 mcg/mL from the initial IVX-411 concentration of 2 mcg/mL in formulation buffer (50 mM Tris, 150 mM NaCl, 0.1 M L-Arginine, and 5% Sucrose, pH 8.0) for measuring the binding with ACE2-Fc receptor, CR3022, and S309, respectively. The diluted samples were blocked by adding one part of casein blocking buffer (Tris-buffered saline with casein and 0.05% Tween-20, Thermo-Fisher) to two parts of diluted test sample and incubated at room temperature for one hour. The blocked samples were serially diluted (1:2 dilutions for measuring adjuvanted IVX-411 binding with ACE2-Fc and CR3022, and 1:1.5 dilutions for measuring adjuvanted IVX-411 binding with S309) in blocking buffer across 96-well PCR plates (Thermo-Fisher). The diluted samples were incubated with an equal volume of test ligand (0.01 mcg/mL of ACE2-Fc or CR3022, or 0.2 mcg/mL S309) overnight at room temperature on a rotator. Ninety-six well polysorp Nunc Immobilizer Amine assay plates (Thermo-Fisher Scientific) were coated with 1 mcg/mL RBD [1] (Institute for Protein Design, Seattle, Washington) [2]diluted in DPBS and incubated overnight at 4°C. The next day the assay plates were washed three times with wash buffer (DPBS with 0.05% Tween-20), blocked with casein blocking buffer for one hr at 25°C, and then washed again as described above. The PCR plates containing the IVX-411 drug product formulations were centrifuged at 1,600 x g for 5 min at ambient temperature and the supernatants were carefully transferred to the assay plates. The plates were then incubated at 25°C for two hrs, washed three times with wash buffer, and incubated with goat anti-human Ig-HRP secondary antibody

Southern Biotech, Birmingham, AL) diluted in blocking buffer (1:5000 dilution for ACE2-Fc and CR3022, and 1:10,000 dilution for S309) for one hr at 25°C. Assay plates were washed three times and developed using TMB substrate solution (Sigma-Aldrich, St. Louis, MO) by incubating at room temperature in the dark for ~15 min and quenched using 1N sulfuric acid. Optical density (OD) at 450 nm was read using SpectraMax iD3 microtiter plate reader (Molecular Devices, San Jose, CA) and the data was analyzed using Origin software (OriginLab, Northampton, MA) after blank subtraction. IVX-411 concentration was calculated from the OD 450 values using parameters obtained from a 4-point logistic fit of the reference sample that was run on each plate.

**Adjuvant adsorption and desorption studies-** Binding of IVX-411 to Alum adjuvants was tested at various IVX-411 concentrations ranging from 50-200 mcg/mL. IVX-411 was diluted to the target antigen concentration in 50mM Tris, 150mM NaCl, 0.1 M L-arginine, 5% sucrose, pH 8.0 along with a final Aluminum concentration of 1.5 mg/mL as Alhydrogel® or 1.5 mg/mL AdjuPhos® (Iniviogen, San Diego, CA). The solution was incubated for 1 hr at room temperature with gentle end over end rotation. Adsorption to the Alum was determined by centrifugation at 4,000 x g for 5 min and analysis of the supernatant by either UV-Vis spectroscopy or SDS-PAGE. Desorption of the antigen was performed by pelleting the Alum with the antigen bound as described above and resuspending the pellet in desorption buffer: 0.4 M sodium phosphate, 1X LDS loading buffer (10% glycerol, 1% lithium dodecyl sulfate (LDS), 0.2 M triethanolamine-Cl pH 7.6, 1% Ficoll®-400, 0.0125% phenol red, 0.0125% coomassie G250, 0.5 mM EDTA, pH 7.5), 50mM DTT and heating at 95°C for 10 min. The solution was then centrifuged to pellet the Alum as described above, followed by analysis by SDS-PAGE on the desorbed antigen in the supernatant. Estimation of the protein concentration by SDS-PAGE was performed by densitometry using ImageJ software (NIH, Bethesda, MD, available at: https://imagej.nih.gov/ij/).

For IVX-411 samples adjuvanted with CpG, CpG was added at a final concentration of 0.3 mg/mL. For CpG and Alum co-formulated samples, IVX-411 was bound first by incubation at room temperature by end over end rotation for one hour as described above. CpG was then added at 0.3 mg/mL and incubated for another hour at room temperature with end over end rotation. The amount of IVX-411 and CpG bound to the Al salt was determined by analysis of the supernatant by SDS-PAGE, and OD260 measurements (using a Nanodrop spectrophotometer (Thermo-Fisher), respectively.

**Langmuir adsorption isotherms-**Langmuir adsorption isotherms were constructed to determine the adsorptive capacity of IVX-411 to AH or AP and to calculate the binding strength of the antigen to the adjuvant. IVX-411 was incubated with a fixed amount of Alum (50 mcg Alum as either AH or AP) as function of increasing protein concentration (0-600 mcg/mL) using 50mM Tris, 150mM NaCl, 0.1 M L-arginine, 5% sucrose, pH 8.0 as the diluent. The mixtures were incubated at room temperature for 1 hr with gentle end over end rotation followed by centrifugation at 4,000 x g for 5 min and the supernatant analyzed by UV-Vis spectroscopy as described above. The protein concentration of the supernatant (Ce) was calculated using the A280 value after light scattering correction of the raw spectra as described above. The data was then fit to the linearized form of the Langmuir equation to calculate the Qmax and KL.

The Langmuir equation is shown below

$$q_e = \frac{Q_{max} K_L C_e}{1 + KL\, C_e}$$

The linearized form of the Langmuir equation:

$$\frac{C_e}{q_e} = \frac{1}{Q_{max}} C_e + \frac{1}{K_L Q_{max}}$$

Where $q_e$ and $C_e$ are solid phase and liquid phase equilibrium concentrations, respectively. $Q_{max}$ is maximum (monolayer) binding capacity. $K_L$ represents the Langmuir isotherm constant, which represents binding strength (the higher the value the stronger the binding.)

**Compounding samples for stability and mouse immunogenicity studies-** IVX-411 bulk drug substance was thawed and diluted to a final protein concentration of 2 mcg/mL in 50mM Tris, 150mM NaCl, 0.1 M L-Arginine, 5% Sucrose, pH 8.0 with either 1) 1.5 mg/mL AH, 2) 1.5 mg/mL AP, 3) 1.5 mg/mL AH and 0.3 mg/mL CpG, 4) 1.5 mg/mL AP and 0.3 mg/mL CpG, and 5) 1X final AV concentration. Samples were diluted in pre-sterilized 2R glass vials (Schott, Mainz, Germany) and stopped with FluorTec® coated pre-sterilized 13mm serum stoppers (West Pharmaceuticals, Exton, PA) followed by sealing with 13mm aluminum seals. All vials were filled in a Class II biosafety cabinet under aseptic conditions. Vials were then stored in at 4, 25, or 37°C until removal the designated stability timepoint. Samples for mouse immunogenicity were shipped overnight on wet ice packs to the Wadsworth Center (Albany, NY).

**Mouse immunogenicity studies-** Mouse immunogenicity studies were performed by the Wadsworth Center under compliance of the Institutional Animal Care and Use Committee (IACUC), as described previously [3]. Mice were immunized subcutaneously with two 50 µl injections of vaccines on study days 0 and 21 injections. Vaccine samples were refrigerated at 4°C between the two doses. Blood was collected from mice via the submandibular vein on study days 21, 35, and 65. Alum and CpG doses were 150 and 30 mcg in 0.1 mL total volume, respectively.

**Pseudovirus Neutralization Assay-** Pseudo-typed lentiviral reporter particles (RVPs) tagged with luciferase (Luc) (Integral Molecular Inc.) were used with 293T-hsACE2 cells (Integral Molecular Inc.) to determine the SARS-CoV-2 neutralizing potential of mouse serum as described previously [3].

**Statistical analysis-** Statistical analysis was performed between immunization groups using Kruskal-Wallis analysis using Dunn's multiple comparisons test using GrapPad Prism software.
